# Seasonal variation in physiology and shell condition of the pteropod *Limacina retroversa* in the Gulf of Maine relative to life cycle and carbonate chemistry

**DOI:** 10.1101/2020.02.25.964478

**Authors:** Amy E. Maas, Gareth L. Lawson, Alexander J. Bergan, Zhaohui A. Wang, Ann M. Tarrant

## Abstract

Natural cycles in the seawater partial pressure of carbon dioxide (CO_2_) in the Gulf of Maine, which vary from ∼250-550 µatm seasonally, provide an opportunity to observe how the life cycle and phenology of the shelled pteropod *Limacina retroversa* responds to changing food, temperature and carbonate chemistry conditions. Distributional, hydrographic, and physiological sampling suggest that pteropod populations are located in the upper portion of the water column (0-150 m) with a maximum abundance above 50 m, allowing them to generally avoid aragonite undersaturation. Gene expression and shell condition measurements show, however, that the population already experiences biomineralization stress in the winter months even when aragonite is slightly oversaturated, reinforcing the usefulness of this organism as a bio-indicator for pelagic ecosystem response to ocean acidification. There appear to be two reproductive events per year with one pulse timed to coincide with the spring bloom, the period with highest respiration rate, fluorescence, and pH, and a second more extended pulse in the late summer and fall. During the fall there is evidence of lipid storage for overwintering, allowing the second generation to survive the period of low food and aragonite saturation state. Based on these observations we predict that in the future pteropods will likely be most vulnerable to changing CO_2_ regionally during the fall reproductive event when CO_2_ concentration already naturally rises and there is the added stress of generating lipid stores.

## 1. Introduction

As human activity increases atmospheric carbon dioxide (CO_2_) levels, dissolution of anthropogenic CO_2_ gas into oceanic waters reduces both pH and the saturation state of calcium carbonate, a process called ocean acidification. Recent studies have suggested that the coastal waters of the US Northeast region, including the Mid-Atlantic Bight and the Gulf of Maine, may be more prone to acidification than previously thought (Wang et al., 2017b; Wang et al., 2013; Wanninkhof et al., 2015). These coastal waters have, on average, lower pH and aragonite saturation states (Ω_Ar_) than other shelf systems, while their buffering capacity is low. This means that they are closer to undersaturation and that pH will change more quickly with the addition of anthropogenic CO_2_.

How the chemistry and biology of the Northeastern U.S. region respond to ocean acidification is of significant importance to the region’s ecosystem health and profitability of wild fisheries, aquaculture, and tourist economies (Cooley & Doney, 2009; Ekstrom et al., 2015; Fay et al., 2017; Hare et al., 2016). Natural variability in the partial pressure of CO_2_ (*p*CO_2_) in the Northeast US coastal shelf is driven primarily by riverine inputs, seasonal cycles in primary production, and changes in sea-surface temperature (Previdi et al., 2009; Salisbury et al., 2009; Wang et al., 2017b). As a consequence, the region already experiences strong natural seasonal variability in *p*CO_2_ levels. The spring season in the Gulf of Maine corresponds to the timing of the lowest levels of surface *p*CO_2_ (∼250 µatm; similar to preindustrial atmospheric levels), while the surface waters reach ∼550 µatm (well over the current global atmospheric average) in the late fall and early winter (Figure 1; Irish et al., 2010; Vandemark et al., 2011; Wang et al., 2017b). The implications of this seasonal variability in *p*CO_2_ on local biota have not yet been examined, but the system experiences a very different type and timing of natural variability compared with what has been documented on the West Coast, which experiences episodic high CO_2_ during the periodic upwelling of CO_2_ enriched subsurface water, with greatest intensity in the spring and summer. Understanding the interactions between this annual variability and organismal responses is important because it could result in periods of predictable high CO_2_ stress that may be mitigated by local acclimation to high CO_2_.

**Figure 1:**
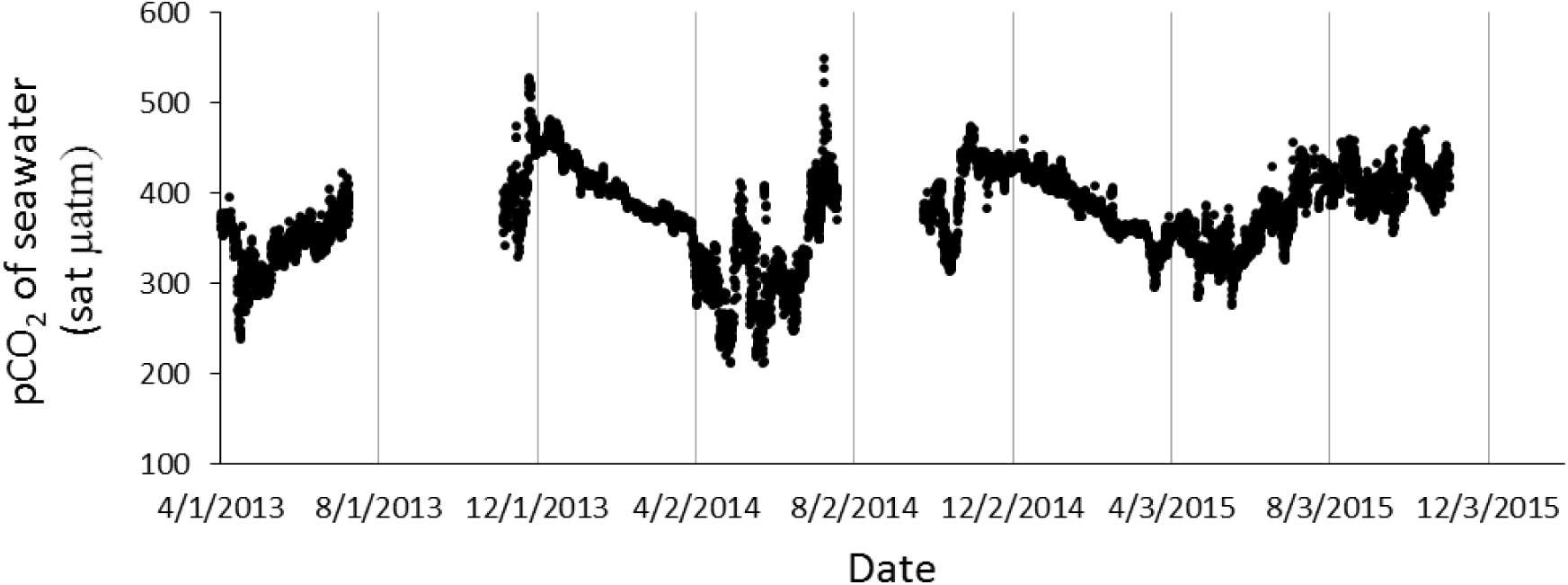
Surface seawater pCO_2_ measured at the Coastal Western Gulf of Maine Mooring (43.02°N, 70.54°W) during the period of this study (Sutton et al., 2014; https://www.pmel.noaa.gov/co2/story/GOM).

The thecosomatous pteropods have been championed as a potentially effective bio-indicator group for studies of ocean acidification stress (Bednaršek et al., 2017; Bednaršek et al., 2012a; Weisberg et al., 2016). These planktonic organisms produce calcium carbonate shells of aragonite that have been clearly shown to be sensitive to aragonite near- and undersaturation in both field and laboratory studies (Bednaršek et al., 2016; Bednaršek et al., 2012b; Bergan et al., 2017; Busch et al., 2014; Comeau et al., 2009; Maas et al., 2018). The species *Limacina retroversa* is a dominant thecosome pteropod in temperate latitudes of the Atlantic, in both the open ocean (Chen & Be, 1964; Wormuth, 1985) as well as continental shelf regions including the North Sea and coastal waters of New England and Eastern Canada (Bigelow, 1924; Kerswill, 1940; Newman & Corey, 1984; Redfield, 1939; Wassmann et al., 1999). The only prior time series of the Gulf of Maine region was conducted in 1933-34 by Redfield (1939), who observed patchy high abundances of small individuals appearing in the fall that persisted through the spring. Redfield noted a correlation between the seasonal progression of the location of *L. retroversa* patches and major surface currents, and based on the bimodal and non-linear growth of their size frequency distributions he suggested that the population is maintained by pulses of lateral advection from an open ocean source. The species has been reported to feed on detritus, diatoms and dinoflagellates (Lalli & Gilmer, 1989; Morton, 1954) with the capacity to ingest large quantities of phytoplankton (∼4000 ng of chl a pigment individual^-1^ day^-1^, Bernard & Froneman, 2009). *Limacina retroversa* is eaten by a variety of fish (Bigelow, 1924; Lebour, 1932; Suca et al., 2018) and can become such a prevalent component of the diet of mackerel and herring that they can act as a vector for toxins and non-palatable compounds, negatively affecting fisheries (Ackman et al., 1972; Foster et al., 2012; White, 1977). They also appear to play a role in modifying the alkalinity budget and carbonate chemistry in deep basins in the Gulf of Maine (Wang et al., 2017b).

The local ecological importance of *L. retroversa*, its natural seasonal exposure to strong variation in CO_2_ levels, and the fact that it is widely distributed in temperate portions of the Atlantic open ocean position the species as a valuable model for the study of East Coast seasonal sensitivity to CO_2_. Previous laboratory studies with this species have documented that exposure to CO_2_ increases their nitrogen excreta (Lischka & Riebesell, 2016). It has also been demonstrated that both the severity and duration of exposure influence respiration rate and gene expression interactively, with increasing duration of CO_2_ exposure widening the difference between the respiratory and gene expression response of individuals exposed to ambient or higher levels of CO_2_ (Maas et al., 2018). These prior experiments, which exposed *L. retroversa* to CO_2_ for 1-14 day durations, indicated that expression of genes associated with biomineralization (particularly chitin, carbonic anhydrase, alkaline phosphatase, collagen, metalloproteinases, mucin, and calcium ion binding) were substantially and increasingly altered when individuals were exposed to longer durations of low (∼460 µatm; Ω_Ar_ = 1.56) versus high (840 µatm, Ω_Ar_ = 0.96 or 1180 µatm, Ω_Ar_ = 0.71) CO_2_. Concurrent to these patterns of gene expression, shell condition consistently showed patterns of decline in response to undersaturation (Maas et al., 2018). These shell condition changes have important ecological consequences, reducing the sinking speeds and mass of the organisms (Bergan et al., 2017; Manno et al., 2012), with potential implications for predator-prey interactions and the efficacy of shell particle fluxes. Juvenile life stages have been shown to be more sensitive to acidification than adults, with increased mortality and a developmental delay during the period of initial calcification when raised at elevated CO_2_ conditions (Thabet et al., 2015). Translating these laboratory experimental results to an understanding of how wild populations experience and respond to a changing marine environment remains difficult. As is the case for most pteropod species, our knowledge of the vertical distribution (which influences their natural CO_2_ exposure), ecology, and in situ physiology of *L. retroversa* in the Gulf of Maine, and the North Atlantic as a whole, remains limited.

The objective of this study was thus to examine the natural in situ distribution and physiology of pteropods in the Gulf of Maine and to determine whether there are changes in the local population that are correlated to seasonal patterns in carbonate chemistry, temperature, and food availability. We hypothesized that annual changes in local carbonate chemistry would have an influence on the metabolic rate, biomineralization and associated gene expression of the local population of *L. retroversa*. Although metabolism and reproduction of other species are known to cycle in temperate/boreal systems in concert with spring blooms and changing temperatures (Bigelow, 1924; Conover & Corner, 1968; Conover, 1959), the annual patterns of development, reproduction and energetic metabolism over annual timescales have not been clearly defined for *L. retroversa*. Understanding these cycles is important because they can reveal periods that could serve as population “bottlenecks” under future anthropogenically acidified conditions. One potential bottleneck is the life stage when initial biomineralization occurs, as it has been demonstrated to be one of the most sensitive for both pteropods and other molluscan groups (Thabet et al., 2015; Waldbusser et al., 2013; Waldbusser et al., 2015; White et al., 2013). It remains mostly unclear when, and due to what physical factors, *L. retroversa* spawns in the North Atlantic. It has been reported based on field sampling that species are capable of producing eggs continuously after reaching a shell diameter of ∼1.1 mm (Hsiao, 1939), while our combined field and laboratory work in the Gulf of Maine has demonstrated that individuals produce eggs during all seasons (Thabet et al., 2015). Thus, characterizing the vertical habitat for these early life stages, and co-locating carbonate chemistry and timing of reproductive events was the second focus of this observational study.

## 2. Materials and Methods

Hydrographic and animal sampling was done concurrently approximately every 3-5 months over a two year period. The gene expression of freshly caught and rapidly preserved organisms was interrogated to provide a snap-shot of in situ physiological activity. Due to methodological constraints, oxygen consumption rate experiments were begun < 24 h after capture and were measured over the course of a 24 h period. Shell condition was measured from the same individuals used in the respiration experiments. Although these measurements were not made immediately after pteropod capture, they represent close to in situ conditions and document real intra-seasonal variation in organismal metrics.

### 2.1 Hydrographic and Animal Sampling

Abundance and distributional sampling of pteropods was conducted in the western Gulf of Maine (GoM) on nine 1-3 day cruises beginning May 22^nd^, August 27^th^, and October 22^nd^ 2013, January 29^th^, April 25^th^, August 19^th^ and November 4^th^ 2014 and April 26^th^ and July 2^nd^ 2015 on the *R/V Tioga.* During each cruise, vertically stratified net and hydrographic sampling was conducted at Murray Basin (42° 21’ N and 69° 47’ W; Figure 2), one of the deepest (ca. 260m) portions of the GoM. A standard 1/4-m^2^ Multiple Opening/Closing Net and Environmental Sensing System (MOCNESS; Wiebe et al., 1985) with 150-µm mesh nets was towed at 8 discrete depths throughout the water column to determine the distribution and density of thecosomes (the group of shelled pteropods that includes *L. retroversa*). All net tows were conducted during daylight hours. The upper nets consistently targeted 0–25, 25–50, 50–75, 75– 100, and 100–150 m. The depths of the lower three nets were chosen adaptively during each cruise based on transmissometer profiles such that the deepest two nets sampled exclusively within the benthic nepheloid layer, in order to examine any associations of pteropods with the particular chemistry of this bottom resuspension zone (see Wang et al. 2017b). Zooplankton samples were preserved in 70% ethanol for later enumeration.

**Figure 2:**
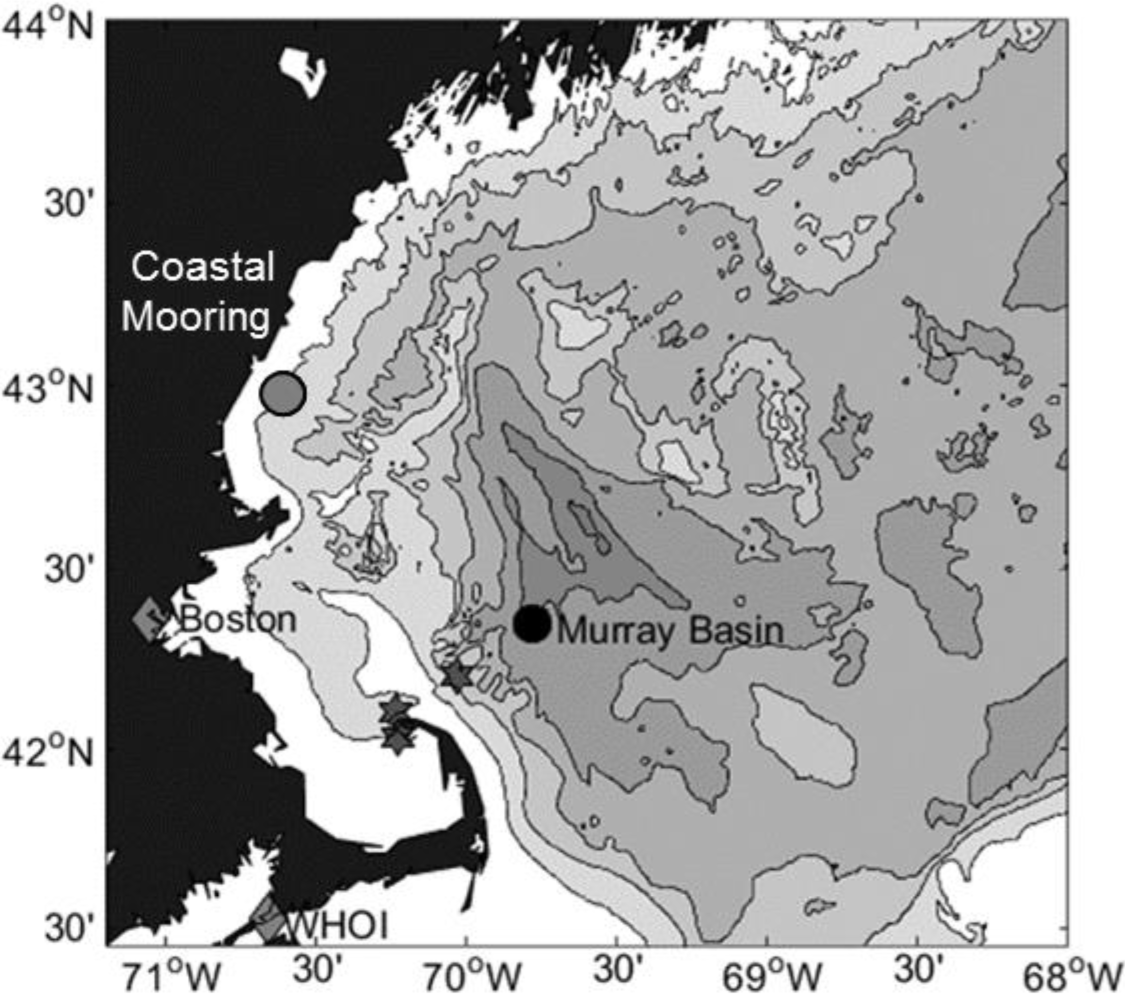
Study region map. Hydrographic and MOCNESS sampling took place at Murray Basin (black circle) during all seasons while animal collection for later physiological analyses took place at multiple additional stations (stars), particularly inshore sites where organisms were most abundant. Position of the Coastal Western Gulf of Maine Mooring is also noted (grey circle).

CTD casts were also routinely conducted at the Murray Basin site using 3-L Niskin bottles and a SBE3/SBR4 sensor set, to characterizing the local hydrography (salinity, temperature, fluorescence, dissolved oxygen, and beam transmission) and carbonate chemistry. Depths for bottle sampling were chosen based on station water depth with a typical profile sampling the upper 100 m depths at 10 m intervals, the 100-200 m depths at 20 m intervals, and less frequently below. Samples for dissolved inorganic carbon (DIC) and total alkalinity (TA) were collected in 250 mL Pyrex borosilicate glass bottles following the best practice of seawater CO_2_ measurements (Dickson et al. 2007). Air head space of about one percent of the bottle volume was left to allow room for expansion. Each sample was then poisoned with 100 µL of saturated mercuric chloride, capped with an Apiezon-L greased stopper, thoroughly mixed, and then tied with a rubber band over the glass stopper.

In the lab, splits of each MOCNESS net sample were examined under a lighted stereomicroscope and *L. retroversa* were enumerated based on size class (<0.5 mm, 0.5-1 mm, 1-3 mm and >3 mm). The relationship between pteropod abundance and hydrographic variables from MOCNESS sampling and available carbonate chemistry data from bottle samples was conducted in PRIMER version 6 (Clarke & Gorley, 2006; Clarke & Warwick, 2001) following the methods detailed in Maas et al. (2014). Briefly, and only using nets where pteropods were present, abundances were square root transformed, and a Bray Curtis resemblance matrix was constructed. Hydrographic data were normalized to a common scale (by subtracting the mean and dividing by the standard deviation), and a BIO-ENV analysis testing all combinations of environmental variables was run with a Spearman rank correlation and permuted 99 times.

For lab measurements of carbonate chemistry, DIC was measured using an Apollo SciTech DIC auto-analyzer, while TA was measured using an Apollo SciTech alkalinity auto-titrator, a Ross combination pH electrode, and a pH meter (ORION 3 Star) based on a modified Gran titration method (detailed in Wang et al., 2017b). pH and aragonite saturation state (Ω_Ar_) were calculated from bottle sample measurements and concurrent temperature and salinity measures from the CTD cast using the CO2SYS program by Pierrot et al. (2006) with constants from Mehrbach (1973) as refit by Dickson and Millero (1987).

During a subset of cruises, adult individuals (>0.5 mm, with fully developed wings/parapodia) were captured for gene expression and respiration experiments. These individuals were collected using a Reeve net with 333-µm mesh and a large cod end that was deployed at slow speeds, for a short duration (< 1 h) with the aim of gently sampling live, undamaged specimens. Although it was originally intended for these organisms to be collected from the deep offshore site concurrent with carbonate chemistry and distributional sampling, as the time series progressed it was found that individuals were unexpectedly more abundant at stations closer to shore. Thus tows were conducted at multiple stations in the Gulf of Maine (C42° 22.1’ - 42° 0.0’ N and 69° 42.6’ - 70° 15.4’ W), and the bulk of live organismal collection occurred at a shallow station close to land (42° 12’ N and 70° 01’ W; Figure 2). *Limacina retroversa* were sorted from other taxa and either placed in locally collected 64-µm filtered refrigerated (8° C) seawater for later respiration experiments, or preserved in RNAlater for transcriptomic analyses.

### 2.2 Respiration Rate

Individuals for respiration experiments were returned via coolers to a walk-in cold-room facility at the Woods Hole Oceanographic Institution within 8 h of collection. The cold-room was set at a constant temperature of 8°C for all experiments to allow for direct comparisons of respiration rates. This temperature was chosen as the average temperature at 50 m depth in the region (it ranged between 6-10°C over our time series). During each experimental period, water had been collected from an offshore site concurrent to pteropod sampling and had been returned to the lab in advance for filtration (0.2-µm) and thermal equilibration. It had been bubbled with ambient air, which ranged from a calculated 380 to 440 µatm over the seasonal cycle, for ∼12 h prior to the arrival of the pteropods. Upon arrival, pteropods were placed in 1-L beakers of water containing the filtered in situ water (15 ind L^-1^) for 8-12 h to allow for further temperature acclimatization and gut clearance. After this period, the healthiest looking individuals (actively swimming or with parapodia extended) were placed into glass respiration chambers (containing optode spots; OXFOIL: PyroScience, Aachen Germany) with 2-3 mL fresh 0.2-µm filtered water. A control chamber, without a pteropod, was set up for every fourth chamber to determine background bacterial respiration. The oxygen concentration was measured non-invasively at the start of the experiment and 24 h later using a FireStingO2 optical oxygen meter (PyroScience, Aachen Germany) as described in Maas et al. (2018). At the conclusion of the experiment, each organism was visually inspected to confirm survival, briefly rinsed with DI water, placed in a pre-weighed aluminum dish, and weighed on a Cahn microbalance (C-33; 1 µg precision) for wet mass. They were then dried for 3-7 days in a drying oven at 60°C and weighed again to obtain dry mass. Final oxygen consumption rates were calculated based on the change in oxygen between the final and initial oxygen measurements, corrected for the bacterial respiration from the control chambers (µmol O_2_ h^-1^).

To test the effect of seasonality on respiration, we first normalized to correct for variations in temperature of the environmental rooms. The rooms achieved 8.1 ± 0.5°C for most experiments, but an equipment failure for the chiller unit during one cruise (August 2014; 5.6°C) resulted in a need temperature correction for cross-comparison. The average temperature of the 24 h respiration experiment was used for the original temperature (T_i_) and the adjusted rates (R_f_) were calculated at 8.0°C using a temperature coefficient (Q_10_) of 2 according to the equation:

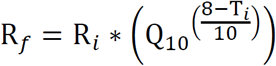

where R_i_ is the final oxygen consumption rate measured for each individual. Although it is known that that Q_10_ is species specific (Seibel et al., 2007), there are no published studies of Q_10_ for *L. retroversa*. Thus, this coefficient value was chosen as it is mid-range for the published Q_10_ of congeners (1.6-2.3; Ikeda, 2014; Maas et al., 2011). Since total variation in temperature across the experiments was small (−2.4 to +0.6 °C), slight variations in actual Q_10_ would not substantially influence the calculated temperature corrected respiration rate. These temperature-normalized respiration rates and the wet mass of each individual were then log transformed to meet the assumptions of linearity for a general linear model. Using the statistical program SPSS, we tested whether wet mass was different among experiments and assessed whether there was an effect of season on the respiration rate of the thecosomes with wet mass as a covariate. Statistical analyses relating hydrographic variables with respiration rate were calculated as Pearson’s correlations, also in SPSS.

### 2.3 Shell Condition

Individuals that had been maintained in captivity for 1-3 days and used in respiration experiments were dried as above, and then analyzed for shell condition following the methods of Bergan et al (2017). Briefly, they were placed in 8.25% hypochlorite bleach for 24-48 h, rinsed in DI water and dried again. These individuals were then photographed under a stereoscopic light microscope at 25x magnification to enable measurements of transparency. Images were analyzed by first identifying the shell against the white background by thresholding the image to black and white. The aperture as well as any holes were manually cropped from the object. The degree of transmittance was then calculated using a custom MATLAB code as the mean greyscale value (range: 0–255) of the pixels of the shell divided by 255 to get a scale of 0 (black) to 1 (white). Data did not meet the assumption of homogeneity of variance so a Welch’s one way ANOVA with Tamhane T2 post-hoc analysis were performed to determine the effect of seasonality.

### 2.4 Gene Expression Analysis

Gene expression analyses were conducted on three pooled RNA samples from each season. These were compared using pairwise differential expression analyses and cluster analysis. Specifically, individuals collected via Reeve net and immediately preserved in RNAlater were frozen at −80°C for less than 12 months prior to RNA extraction. Total RNA was extracted from pooled samples of 5-6 individuals using the E.Z.N.A. Mollusc RNA Kit (Omega Biotek). RNA yield and purity were assessed using a Nanodrop spectrophotometer. Three RNA samples were then selected for each of the four seasonal cruises based on yield and purity (12 total samples), and these were submitted to the University of Rochester Genomics Research Center for RNA quality assessment (Bioanalyzer), library construction and sequencing. Libraries were constructed using TruSeq Reagents and then sequenced on 1 lane of an Illumina HiSeq 2500 as a High Output v4 125 bp PE project. The sequencing facility used Trimmomatic (v.0.32; Bolger et al., 2014) to eliminate adapter sequences (2:30:10) and removed low quality scores using a sliding window (4:20), trimming both trailing and leading sequences (13) and leaving only sequences with a minimum length of 25 for downstream use.

Reads from individual RNA samples were then aligned to a previously assembled and annotated *L. retroversa* transcriptome (Maas et al. 2018; GBXC01000000) using Bowtie2 (v.2.2.3; Langmead & Salzberg, 2012). This transcriptome is sufficiently complete (BUSCO score C:90.6% [S:65.1%, D:25.5%], F:8.0%, M:1.4%, n:978; Simão et al., 2015) to support the analysis. Estimates of abundances were made with RSEM (Li & Dewey, 2011), and a pairwise comparison of differential expression (DE) between seasonal expression profiles was performed using an edgeR analysis with R v.3.0.1 (Robinson et al., 2010), and relying on the pipeline packaged with Trinity (v.2.1.1; Haas et al., 2013). Genes were defined as DE if the log_2_-fold change was > 2 (corresponding to a four-fold change in expression), and using a false discovery rate and p value < 0.05. The BLASTX and Gene Ontology (GO) annotations for all DE genes were retrieved from the previous annotation, and a GO enrichment analyses was conducted with Blast2GO (Conesa et al., 2005), applying a Fisher’s Exact Test with a p-value and false discovery rate (Benjamini and Hochberg) cutoff of < 0.05 for each seasonal comparison. Results were reported using the most specific GO term available. Total patterns of gene expression were compared using non-metric multidimensional scaling (nMDS) and ANOSIM using Primer (v. 7; Clarke & Gorley). TMM normalized FPKM values were plotted as an nMDS plot using a Bray-Curtis similarity matrix.

A priori we hypothesized that there would be seasonal differences in expression of genes associated with CO_2_ exposure, biomineralization, development, reproduction and energetic metabolism. To test these hypotheses, we first identified genes that were previously demonstrated to be responsive to CO_2_ in laboratory exposures of the same species (Maas et al. 2018). We also searched for transcripts that have been identified as being associated with the mantle and biomineralization in Mollusca by assembling a BLAST database using the GenBank accession numbers provided in Zhang et al. (2012). Following the same procedure we searched for genes previously identified as being associated with molluscan development and reproduction (Boutet et al., 2008; Ciocan et al., 2011; Tong et al., 2015). We then conducted a TBLASTN search (e-value 1e-5) of these databases and did a reciprocal BLASTX search (e-value 1e-5) of our *L. retroversa* assembly to find putative homologs within our DE transcripts (Supplementary Data 1). To investigate energetic metabolism, all transcripts annotated with the GO terms “metabolic process” or, due to our interest in overwintering, containing the term “lipid” were identified. Heat maps were plotted of each of these subsets of genes using the R package “heatmap.plus”, using the default dendrogram algorithm to determine whether there was significant clustering of the samples based on season.

## 3. Results

### 3.1 Hydrography and Distribution

Hydrographic data were consistent with classical patterns of winter mixing followed by a spring bloom and summer stratification, with lowest pH and aragonite saturation state during the winter and highest during the bloom periods. Specifically, in Murray Basin (∼260 m depth), there was a very cold (∼6 °C) and deeply mixed winter water column. During the spring the surface waters began to warm (∼10 °C) and stratify, supporting a pronounced spring bloom which occurred around May as indicated by fluorescence measurements (Figure 3); below 30 m, however, the water column remained cold. Throughout the summer and fall the surface became increasingly warm (> 18 °C) while some portion of the mid-water (60-180 m) remained at winter temperatures (∼6 °C). The lowest aragonite saturation state in the upper water column (0-60 m) was observed during the single winter sampling event (January 2014), when high DIC and TA resulted in an upper water column value of Ω_Ar_ ∼1.5. During the spring the saturation state returned to typical levels (Ω_Ar_ ∼1.8 in the top 60 m), with peaks in pH (> 8.1) during the spring blooms. In the spring and summer months carbonate chemistry changes were driven both by biological (photosynthesis and respiration) and physical processes (stratification), while a spring freshening contributed to spatial variability in carbonate chemistry (detailed in Wang et al., 2017b).

**Figure 3:**
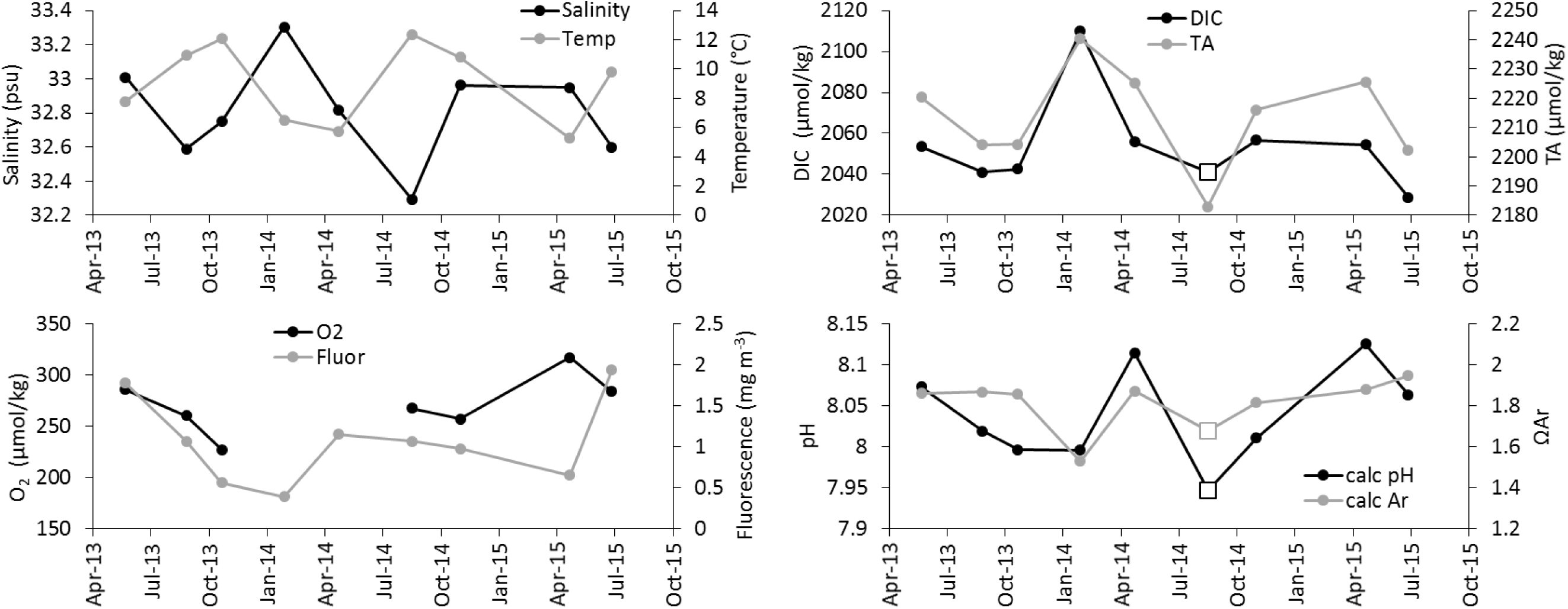
Hydrographic and carbonate chemistry measurements within Murray Basin in the Gulf of Maine (42° 21’ N; 69° 47’ W) over the course of seasonal sampling. Salinity, temperature, oxygen, and fluorescence are calculated as the average values from 0-60 m of the upcast of the CTD. The oxygen sensor was broken for two cruises, resulting in a gap in sampling. Dissolved inorganic carbon (DIC) and total alkalinity (TA) are the average of the measured bottle samples collected from 0-60 m, with one missing DIC value in summer 2014 that was estimated using August 2013 DIC measurements (white box). pH and aragonite saturation state (Ω_Ar_) are the average calculated values from 0-60 m based on measured temperature, salinity, DIC and TA. Those that were calculated using the estimated DIC are denoted with a white box.

Although the upper water column (< 50 m) was clearly the preferred habitat of the thecosome pteropods, there was often a substantially smaller subsurface peak deeper in the water column (100-120 m; Figure 4). In winter, when the water column was more homogeneous and individuals tended to fall in the middle size classes, this second peak was of more similar density to the surface peak. In almost all seasons individuals were completely absent below 200 m. The only exception to this was during July 2015 during a massive bloom of individuals where a few individuals were found all the way to the bottom. Analysis of the size class component of the daytime vertical distribution shows that individuals of the smallest size were significantly more abundant and were preferentially found in the top 50 m of the water column, while those in the 0.5-1 mm size class were spread more evenly through the top 150 m (Figure 5A). The 1-3 mm size class appeared to increase slightly in abundance with depth to 150 m. In contrast, the relatively rare largest individuals (> 3mm) were only ever found in the top 50 m.

**Figure 4:**
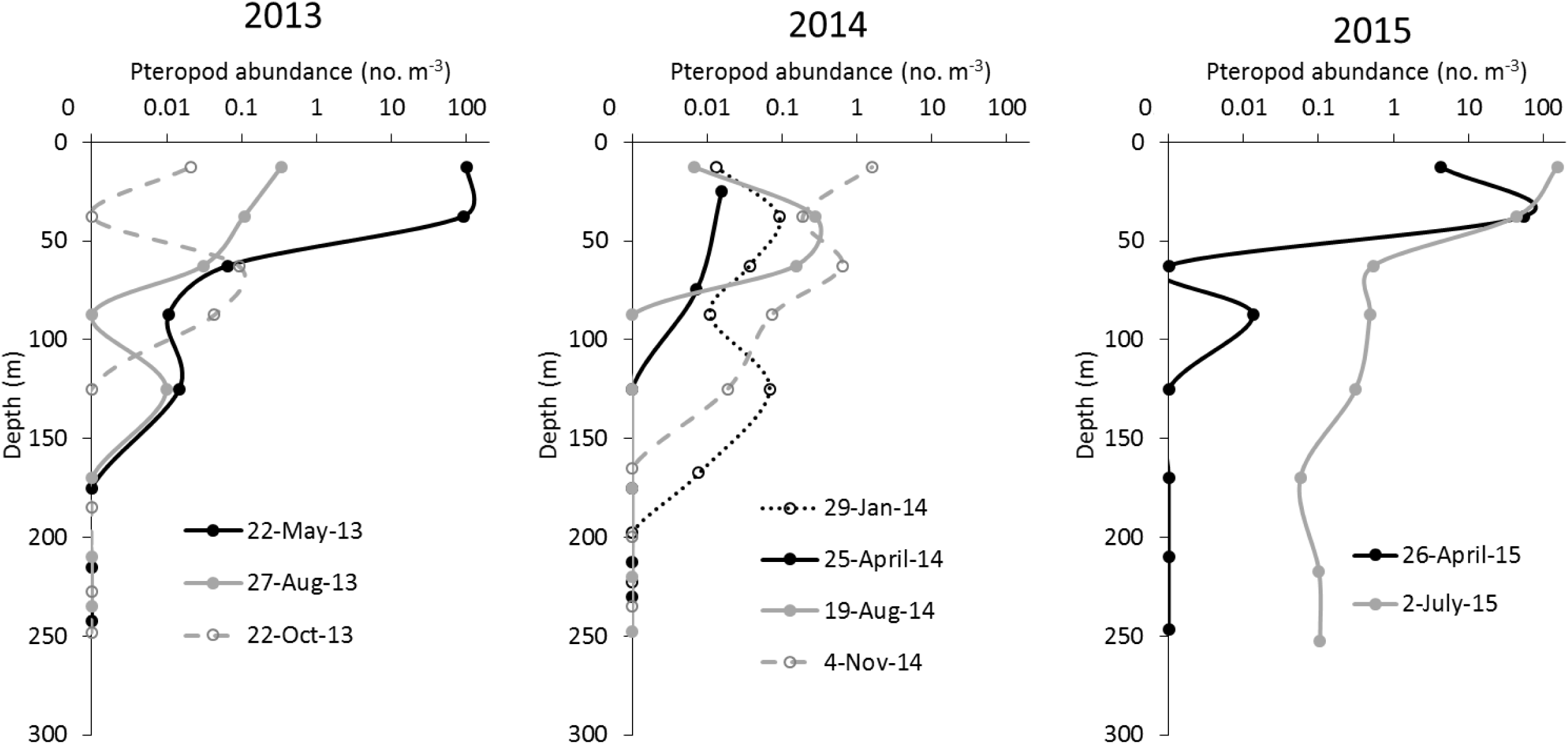
Seasonal vertical distribution of *Limacina retroversa* during 3 years of sampling. The total abundance of *L. retroversa* (all sizes combined) in the water column. The abundance is plotted on a log scale and is respective to the middle of each sampled depth interval.

**Figure 5:**
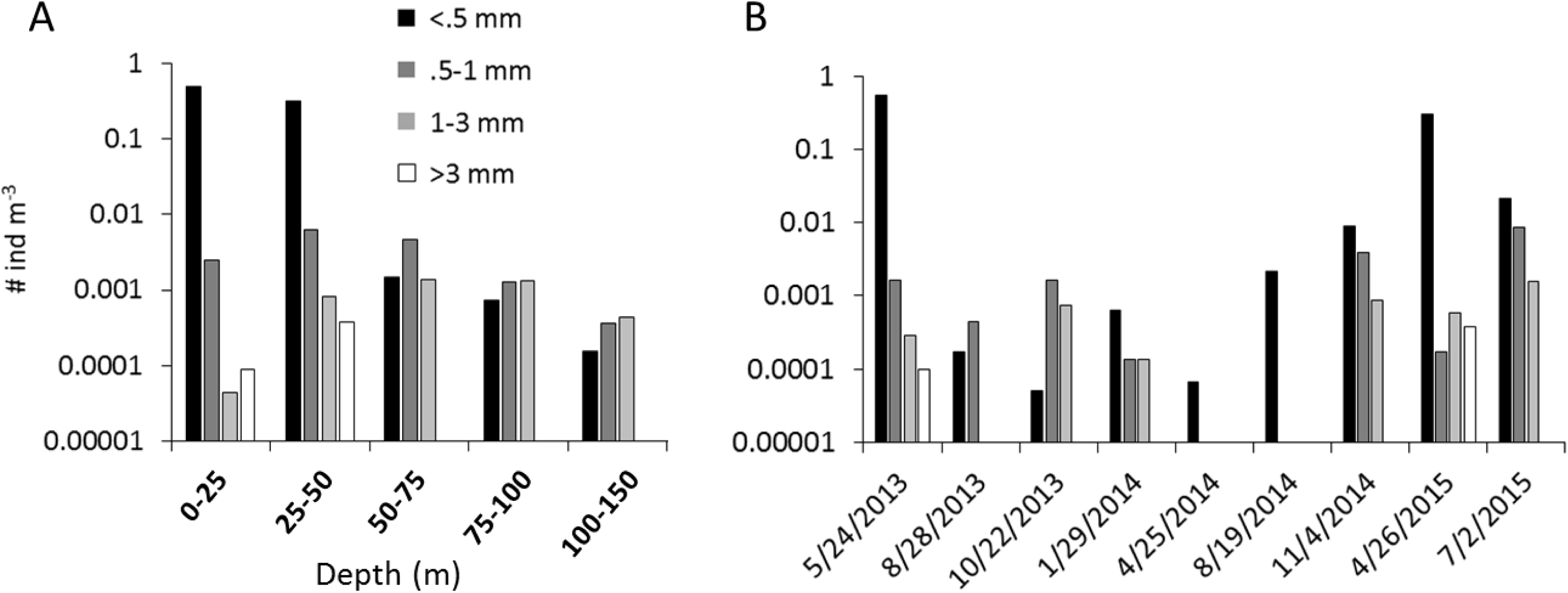
(A) Average vertical distribution of *L. retroversa* in the upper water column (0-150 m) for all seasons combined, and (B) total seasonal abundance by size class for individuals collected from the Murray Basin site in the Gulf of Maine from the full May 2013-July 2015 time-series. Note the logarithmic scale.

During the period of the spring bloom and early summer (April – July) the pteropods were typically found in substantially greater numbers. During this season in 2013 and 2015 was the only time when and individuals of the largest size class (> 3 mm), were present (Figure 5B). In spring 2014, however, sampling occurred in a region of lower pteropod abundance, and no large organisms (> 0.5 mm) were found. In the late summer and fall (August-November) animals began to shift lower in the water column. Fluorescence was the best single predictor of pteropod abundance (Table 1; R=0.325; n=39; p<0.01) as assessed using a BIO-ENV that compared a Bray Curtis resemblance matrix of pteropod abundance (using only nets where pteropods were present) to available hydrographic information from MOCNESS net tows (salinity, temperature, fluorescence, and pressure averaged over the depth interval of the net sample), and bottle data of carbonate chemistry (DIC, TA, and calculated pH, ΩAr, and pCO_2_; corrected for missing carbonate chemistry samples using linear interpolation (n=7)). When pteropod abundance was similarly analyzed using size class information to generate the similarity matrix, pH was the best single predictor (Table 1; R=0.300; n=39; p<0.01).

**Table 1:** Best predictors of pteropod abundance and organismal level metrics. Pteropod abundance, respiration rate, and shell condition were each correlated with available environmental variables. The best single predictor is reported along with its R value, number of measurements and p-value.

| Pteropod metric | best single predictor |  |
| --- | --- | --- |
| Total pteropod abundance | fluorescence | $R=0.325$; $n=39$; $p<0.01$ |
| Size-specific pteropod abundance | pH | $R=0.300$; $n=39$; $p<0.01$ |
| Respiration | fluorescence | $R=0.654$; $n=5$; $p = 0.231$ |
| Respiration excluding Jan | fluorescence | $R=0.8572$; $n=4$; $p = 0.058$ |
| Shell condition | temperature | $R=-0.622$; $n=5$; $p=0.263$ |

### 3.2 Respiration experiments

The temperature corrected respiration rate (µmol O_2_ h^-1^) of *L. retroversa* was significantly different between seasons of sampling with mass as a covariate (F_4, 42_ = 0.176, p < 0.001). Transformed data met the assumptions of homogeneity of variance and normality, and a Bonferoni post-hoc test indicated that spring metabolic rates were substantially higher than rates measured during the fall. This was clearly not a function of changes in organismal size over the seasons as mass varied independently from mass corrected metabolic rate (Figure 6). Of the environmental variables available for all cruises (salinity, temperature, fluorescence and TA), respiration rate was best correlated with fluorescence, but this relationship was not statistically significant (Table 1; Pearson’s 2-tailed correlation; R=0.654; n=5; p=0.231).

**Figure 6:**
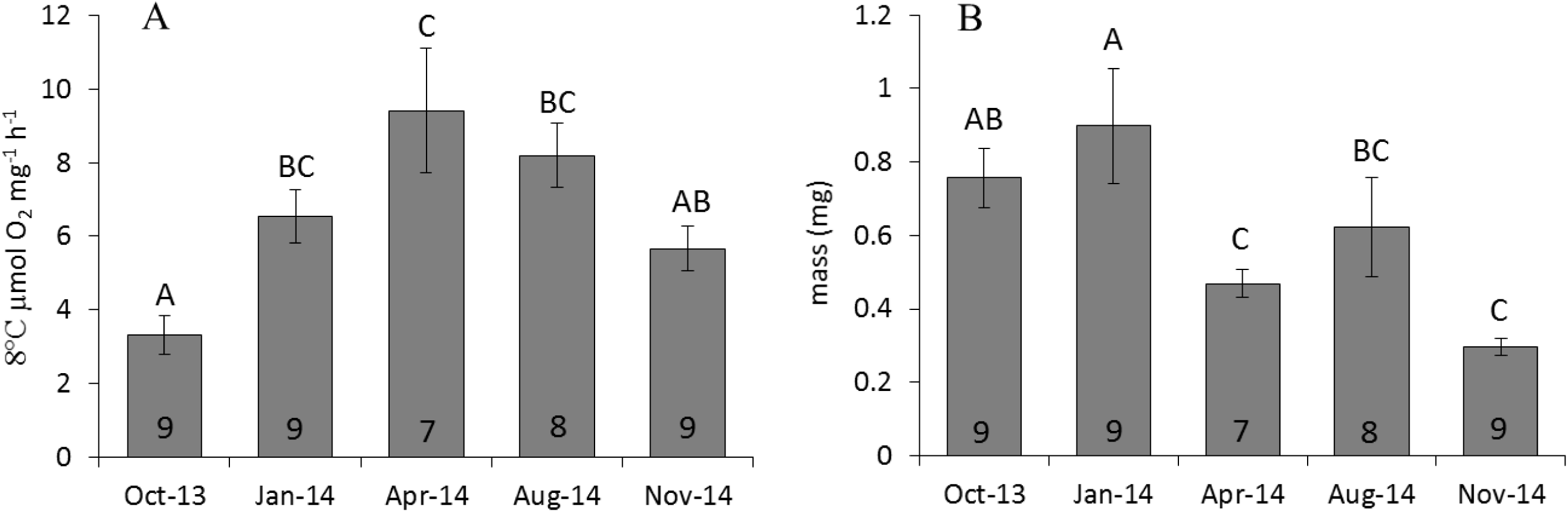
(A) Temperature and mass corrected respiration rates (mean ± SE) of *Limacina retroversa*. (B) Organismal mass (mean ± SE) from the same individuals. The number of individuals per analysis is noted at the bottom of each bar and letters show significant differences among seasons p< 0.05. It is important to note that many of these individuals were collected from nearshore sites so their size distribution is different from that reported in Figure 4.

### 3.3 Shell condition

There was a significant difference in shell transparency across seasons for individuals kept in captivity for up to 3 days (Welch’s F_4, 23.944_ = 79.908, p < 0.001; Figure 7). Based on a Tamhane T2 post hoc test, the lowest transmittance shells were from January and April of 2014, while the highest condition were in August 2014 and April of 2015. Shell condition was best correlated with temperature, but this relationship was not statistically significant (Table 1; Pearson’s 2-tailed correlation; R=-0.622; n=5; p=0.263).

**Figure 7:**
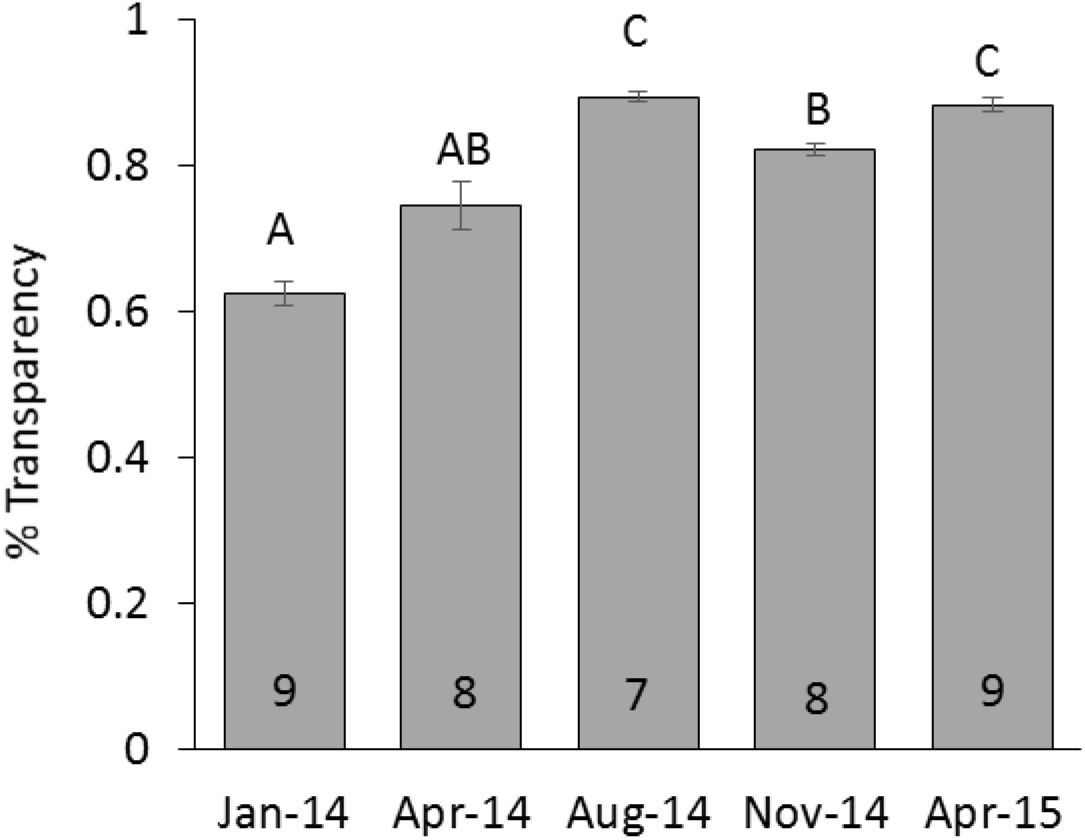
The transparency of *Limacina retroversa* across the seasonal analysis (mean ± SE). Analyses of dried shells from individuals kept in captivity for ∼3 days indicate that the winter individuals had the lowest transparency, while those in the summer individuals had the highest (letters show significant differences among seasons p < 0.05). Spring seasons were distinctly different between years, with 2014 being substantially closer to winter transparency and 2015 values being closer to summer values. The number of individuals per analysis is noted at the bottom of the bar.

### 3.4 Gene Expression Analysis

Comparison of the similarity of transcriptomic profiles across seasons revealed that samples clustered significantly by season (ANOSIM; R=0.985; p = .001) on an nMDS (Figure 8). Non-hierarchical (kR) clustering with progressively larger a priori cluster numbers indicates that the first split (two clusters; ANOSIM: R=0.928; p = .002) places January and April in one group in contrast to November and August. Adding a third cluster does not substantially increase the fit (ANOSIM: R=0.930) compared to two clusters, and was well below the fit for four, which was identical to the seasonality ANOSIM.

**Figure 8:**
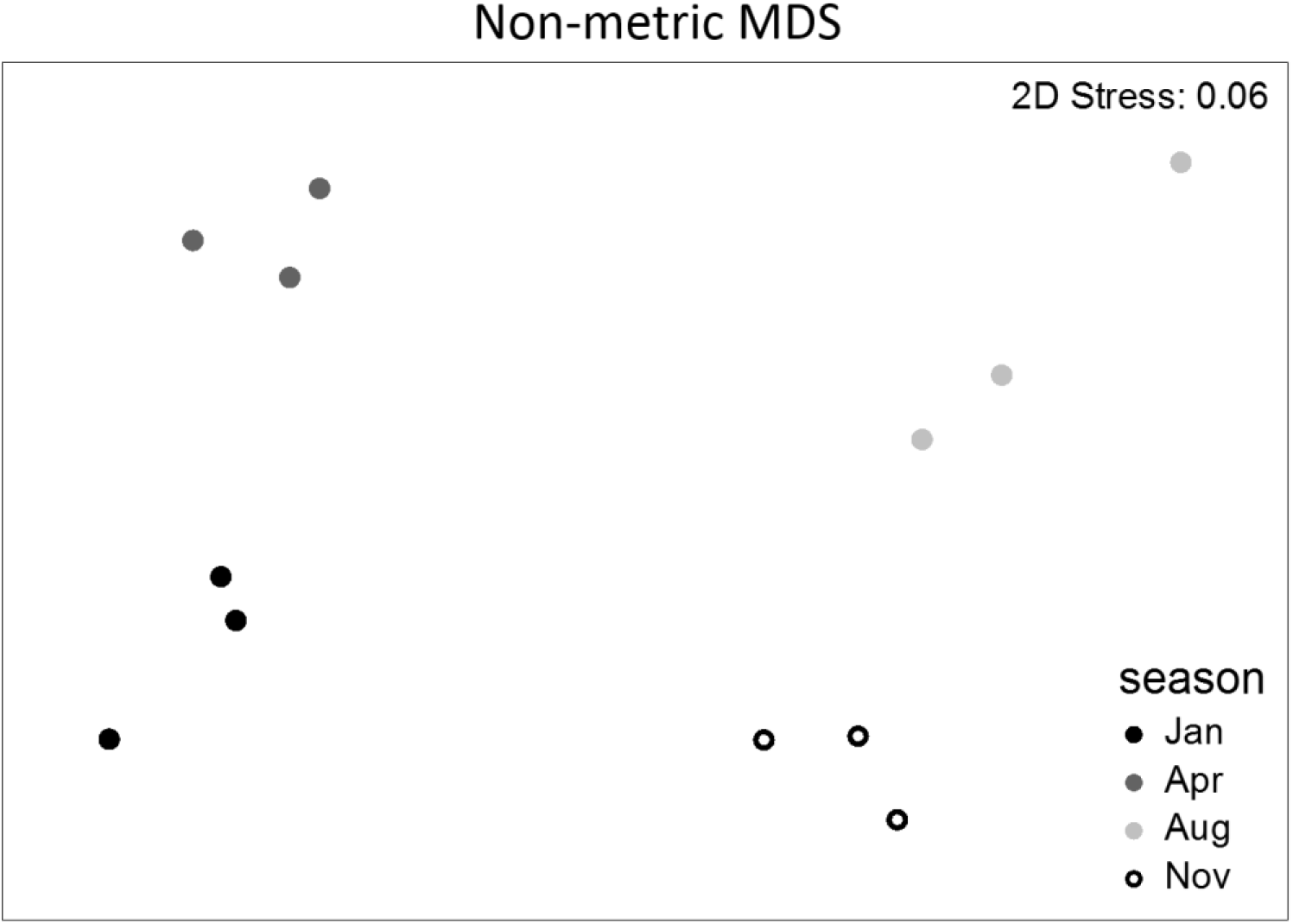
Statistical grouping of samples based on the similarity of their total gene expression profile. The plot is a non-metric Multi-Dimensional Scaling (nMDS) analysis of total TMM-normalized FPKM gene expression based on a Bray-Curtis similarity matrix. Clusters, which are annotated by season (color), are significant (R= 0.985, p < 0.01) based on an ANOSIM.

On average, 68.60% of all sequences from the samples mapped back uniquely onto the *L. retroversa* assembly (Supplementary Data 2). In pairwise comparisons, a total of 4391 differentially expressed genes were identified from four seasonal sampling trips (Figure 9; Supplementary Data 3). This constitutes ∼3.7% of the transcriptome. Although there were distinct differences among the seasons, January and April tended to cluster together in expression pattern as did August and November. The biggest number of DE transcripts was observed between January and August (2324 DE genes) with 740-1411 DE genes identified in other seasonal contrasts. GO enrichment analysis suggested enrichment of lipid transport, oxidoreductase, ATPase, and hydrolase activity in August and November in pairwise contrasts to January and April (Supplementary Data 4). All seasons showed enrichment of chitin binding and chitin metabolism GO terms in comparison with August.

**Fig 9:**
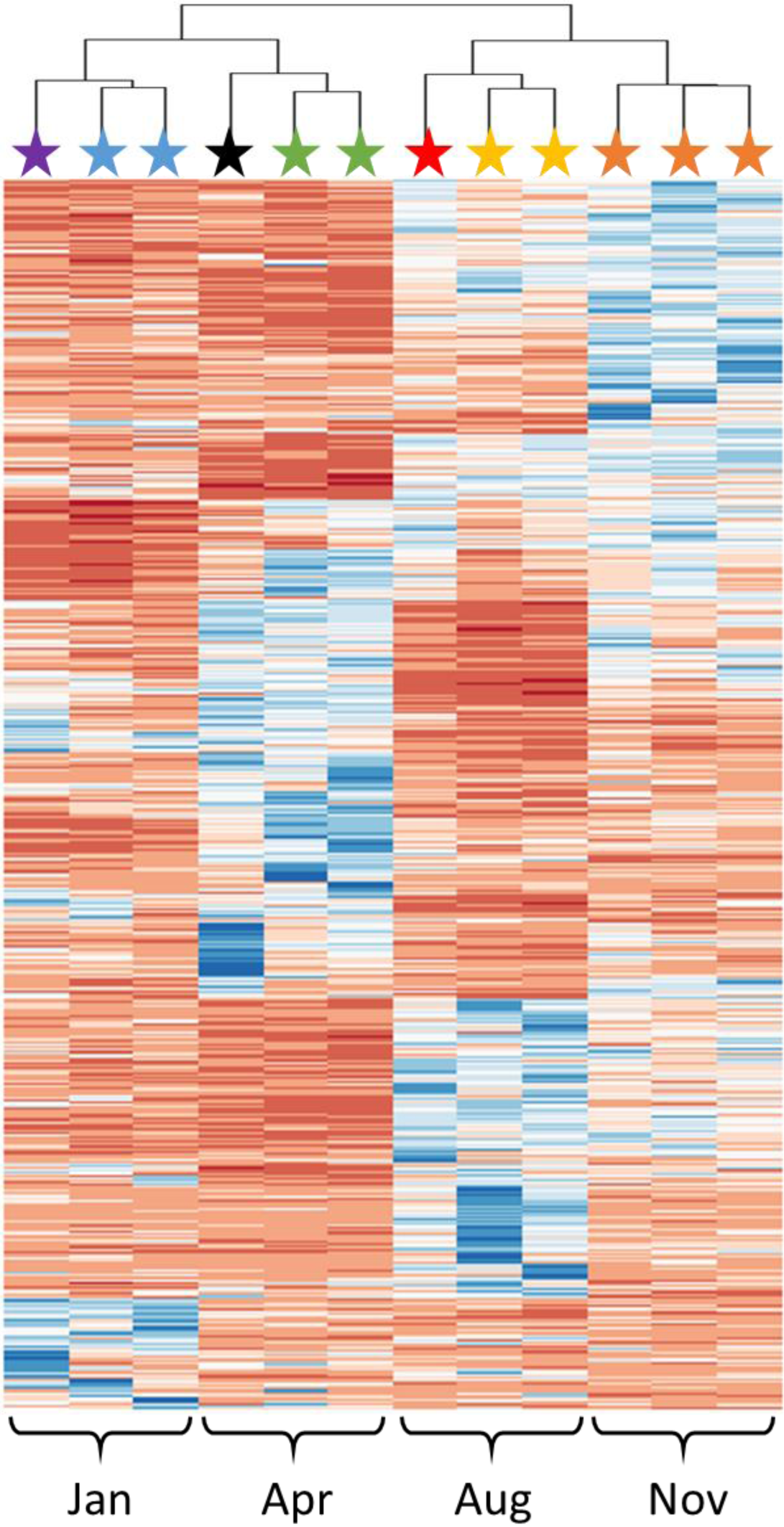
Dendrogram clustering of sample similarity based on expression of the DE subset of genes (TMM normalized FPKM values for DE genes). Each gene was normalized ((x-xmin)/(xmax-xmin)) and a dendrogram was created using the heatmap.plus function in R (Blue = value of 0, Red = value of 1). Clusters were tested for statistical significance (p < 0.05, colored stars) using the Primer v7 statistical package using a Simprof test with 999 permutations.

There were 310 transcripts present in our seasonal DE analysis that had previously been identified (Maas et al. 2018) as being DE in *L. retroversa* after laboratory exposure to low (∼460 µatm; Ω_Ar_ = 1.56) versus high (840 µatm, Ω_Ar_ = 0.96 or 1180 µatm, Ω_Ar_ = 0.71) *p*CO_2_. Analysis of this subset of DE genes indicated that seasons generally clustered, with January separate from all other seasons (Supplementary Data 5). During this period, when in situ saturation state was lowest (Ω_Ar_ = 1.5 compared to Ω_Ar_ = 1.8), the DE patterns in biomineralization (alkaline phosphatase, mucin, calponin) and the extracellular matrix (galectin, plasminogen, fibrocystin) were more similar to the expression patterns of high versus low level laboratory exposure to CO_2_. Focusing on the transcripts identified as being associated with biomineralization as annotated by Zhang et al. (2012), heatmaps of expression showed seasonal clustering (Supplementary Data 5), and 11% (157/1447) were DE in our seasonal analysis. These included shell matrix, chitin, collagen, and c-type lectin transcripts that were upregulated in August and November. Other biomineralization associated transcripts, including alkaline phosphatase, carbonic anhydrase precursor, nacrein, aragonite protein precursor, serine proteinase inhibitor and tyrosinase transcripts were, however, downregulated in August and upregulated in January.

Of the transcripts that were identified by reciprocal BLAST searches as associated with reproduction and development (Boutet et al., 2008; Ciocan et al., 2011; Tong et al., 2015), only 4% (110/2523) were DE in our seasonal analysis. A dendrogram of this subset of genes was clustered by season, but again lumped January and April in strong contrast to August and November (Supplementary Data 5). Transcript expression in April, and to a lesser extent January, were suggestive of late testes development with upregulation of senescence-associated protein, vitelline coat lysin and downregulation of beta-tubulin and testis-specific serine/threonine kinase. In contrast, transcripts associated with egg production (vitellogenin), germline development and sex differentiation were upregulated in August and November. Dendrograms of transcripts annotated as “metabolic process” clustered by season, while those annotated as relating to “lipid” clustered January and April together as separate from strong August and November clusters. Differentially expressed genes suggest that there is strong lipid storage in the second half of the year with genes related to lipoprotein synthesis and transport, several related to lipid synthesis, and lipid metabolism showing significant upregulation in contrast to January and April. This is consistent with GO enrichment analyses, as discussed below.

## 4. Discussion

Our environmental sampling, paired with organismal, distributional and gene expression analysis shows that there is some seasonal exposure to reduced aragonite saturation state in the winter (Ω_Ar_ = 1.5 compared with 1.8 to 1.9) that appears to significantly influence shell condition and the expression of associated biomineralization genes. Reproductive development and changes in lipid metabolism over seasonal cycles were also reflected in our gene expression data, and these processes may, in the future, interact with changing CO_2_ conditions to influence the seasonal sensitivity of *L. retroversa* in the region.

### 4.1 Vertical and Seasonal Distribution

A major contribution of this work is the characterization of the seasonal and size-segregated vertical distribution of *L. retroversa*, which revealed a consistent high peak of small individuals in the upper water column. Throughout the year the peak in abundance tended to correspond with the area of highest fluorescence (0-60 m; often with a max near 30m), while there was often a small deeper population, generally between 100-120 m, containing larger individuals. Statistical analyses of correlation validated this general trend, indicating that fluorescence was the best predictor of total abundance (which is dominated by the smallest size class). This analysis of vertical distribution is consistent with previous estimates of *L. retroversa* vertical distribution with a peak at 50 m and a range of 0-120 m (Bigelow, 1924; Newman & Corey, 1984; Wormuth, 1981). The middle size classes of animals (0.5-3 mm) tended to be slightly deeper in the water column, while the smallest and largest size classes were more likely to be found in the top 50 m, perhaps indicating ontogenetic variation in habitat. When size class is considered, pH becomes the best predictor of abundance, possibly due to the exclusive presence of large individuals during the spring bloom – a period of distinctly higher pH. As phytoplankton abundance (i.e. food availability), pH and temperature are correlated, determining the driver of distributional patterns is not possible. The correlations do, however, emphasize that abundance of various size classes and vertical habitat appear to have a clear seasonality.

The springtime presence of many large and small individuals, as well as examinations of the reproductive tracts of *L. retroversa* in April and May, previously suggested that the early spring is the time of spawning for this animal in the GoM (Hsiao, 1939; Redfield, 1939). Work by Chen and Be (1964), in the subarctic Atlantic on *L. retroversa* suggested a spring spawning, but also continuous reproduction from May – October and a cessation of reproduction during overwintering. In the Bay of Fundy, Canada, two pulses of juveniles were documented in July and November, with reproductive hermaphrodites dominating and likely reproducing in March and August. In the Eastern Atlantic, off of Plymouth, England, *L. retroversa* is reported to continuously spawn through the spring and summer, although there is no information about reproduction during the rest of the year (Lebour, 1932). The findings of these earlier researchers were corroborated in the GoM by observations of egg laying in the lab after seasonal sampling (Thabet et al., 2015), which documented egg production in all seasons but noted that it was particularly high in April and October.

Consistent with these previous studies, the springs of 2013 and 2015 were the periods with the highest number of very small individuals (< 0.5 mm). Based on their size they are likely a new age cohort of sexually undifferentiated individuals (Hsiao, 1939). These two spring time points were also the only seasons when the largest individuals (> 3 mm) were captured. Due to the large number of small animals, the average size of the organisms was lowest in the spring. There was, however, a peak in the second largest size class (1-3 mm; hermaphroditic reproductive adults) and a secondary peak in the smallest size class (< 0.5 mm) in the late fall and early winter 2013-2014. These findings are consistent with a first cohort that grew rapidly through the late fall, consistent with the temperature-size rule for development (Atkinson, 1994). The second cohort would overwinter, growing slowly at the colder winter temperatures and reach the largest average size observed in our study (January) and then reproduce in the early spring. Size distributions from the respiration experiments, which were collected more often from the near-shore sites, also seem to show two generations per year, with smaller, younger individuals in the spring and late fall. A similar reproductive pattern has been proposed for *Limacina helicina* and for other more northern and southern populations of *Limacina retroversa* (Dadon & Cidre, 1992; Newman & Corey, 1984; Wang et al., 2017a).

The timing of the peak abundance of *L. retroversa*, and the information on vertical distributions, suggest that reproduction occurs during periods with high food availability and, during the spring bloom, high pH. The saturation state is relatively stable throughout the spring and summer when the more sensitive juveniles (Thabet et al., 2015) are present. The only season when the pteropod population experiences lower saturation states (winter) is likely the period with lowest reproduction (as suggested by a lower number of small individuals), incidentally protecting this sensitive life stage from the naturally lower saturation state.

Although peaks in pteropod abundance were typically observed in the late spring and early summer, there was high inter-annual variability at our sampling location. In 2014 we did not observe a spring peak at Murray Basin, but we found higher concentrations of larger pteropods at other sites closer to shore that were not sampled quantitatively or in a depth-stratified manner. This is consistent with previous studies of *L. retroversa* in the region which showed patchy distributions and the presence of individuals throughout the year (Bigelow, 1924; Redfield, 1939). This patchiness of zooplankton in the Gulf of Maine is not unique to pteropods, as previous work on the dominant macrozooplankton, the copepod *Calanus finmarchicus*, demonstrated variations in abundance of three orders of magnitude in the same region during the spring bloom with profound implications for transfer of energy to higher trophic levels (Wishner et al., 1988). Studies of various copepod species in the region suggest that these aggregations are due to both physical factors (such as fronts, downwelling zones, or advection), as well as biological forces (development time, bloom timing, species interactions, (Ji et al., 2009; Wishner et al., 1995).

In addition to reproductive pulses and general patchiness, it is likely that we are observing multiple cohorts of *L. retroversa* that are being advected onto the shelf from offshore with size not exclusively representative of local reproductive timing. Bigelow (1924) reported *L. retroversa* as one of the endemic pelagic species of the GoM, speculating that the population sustains itself with local reproduction. Redfield’s work, however, indicated that this species’ presence in the system might be maintained by large influxes of immigrants advected into the Gulf from off-shelf regions, rather than through endemic production (Redfield, 1939). Our results support this finding, with high variability in the abundance of the smaller juvenile size classes. Gene expression results do, however, imply that there are distinct changes in reproduction associated transcripts over the course of the study, with a January/April versus August/November clustering. This may suggest a smeared pattern of reproduction with individuals being introduced from offshore from upstream reproduction overlaid on local reproduction coinciding with phytoplankton blooms. The methodology of the gene expression analysis, which pooled multiple individuals, may have contributed to the observed pattern of reproductive genes as large older adults may have dominated the RNA pool, and be more characteristic of the distinct adult age cohorts rather than a representative picture of the dominant reproductive and developmental transcriptome of the community. The strong evidence of two distinct clusters in gene expression pattern strengthens the argument for two major generations per year with differing reproductive strategies, and suggests that there is some plasticity in the phenology of reproduction that is supported by these immigrants.

The presence of these offshore recruits is important because it suggests that the adaptation or acclimation of pteropods to environmental conditions will be in the context of exposure on a scale larger than just the Gulf of Maine. Thus, although pteropods in the Gulf of Maine experience a shelf system that already shows seasonal variation and sensitivity to acidification (Wang et al., 2017b; Wang et al., 2013) it remains unclear whether they could be locally adapted, since sub-populations may be coming from offshore regions with no prior exposure. However, upstream areas where sub-populations may originate are also acidifying, making a broader understanding of the spatial and temporal variability in the carbonate chemistry of the Northeastern U.S. and Canadian coastal waters, as well as Northwestern Atlantic open ocean conditions, of greater relevance for our understanding of the GoM ecosystem’s response to OA. Population analyses using SNP-based methods (such as RAD-seq, as in Blanco-Bercial & Bucklin, 2016), have a better chance of establishing what proportion of individuals of *L. retroversa* in the GOM come from these various regions. As we increasingly work to understand how ecosystems will evolve under anthropogenic change, assessments of population-level genetic exchange, exposure, and sensitivity will become increasingly important for management assessments.

### 4.2 Cycles in Energetic Investment

Characterizing the change in zooplankton respiration over seasonal cycles is important as it has previously been demonstrated to be responsive to a number of factors beyond organismal size and experimental temperature, including food availability (Conover, 1959; Hochachka & Somero, 2002; Ikeda, 1977; Maas et al., 2011). This has been clearly documented in copepods, but data on other groups is scarce. Understanding these linkages is important as zooplankton resilience to stressors, including ocean acidification, has been shown to be contingent on food availability (Pansch et al., 2014; Seibel et al., 2012; Thomsen et al., 2013). Our respiration experiments, which were all conducted at a similar temperature, document a higher mass specific metabolic rate during April, likely due to increased food availability associated with the spring bloom. Regressions of metabolic rate with fluorescence suggest a good correlation during the period of stratification (April – November; R^2^ = 0.8572, p = 0.058), but the correlation loses fit when the January time point is added (R^2^ = 0.654, p = 0.231). This may be because during the very well mixed and low light winter pteropods depend heavily upon non-fluorescent prey (Lalli & Gilmer, 1989). There were no strong patterns of gene expression indicative of changes in aerobic respiration, which may indicate that variation in respiration rate is controlled beyond the transcriptional level.

There were strong transcriptomic signals of seasonal influences on other metabolic processes, with lipid transport, oxidoreductase, ATPase, and hydrolase showing significantly higher representation in the GO terms in the fall (GO enrichment; August/November) when compared with the earlier time points. Directed investigation of lipid associated terms resulted in a similar clustering of samples, suggesting that there was greater lipid transport and lysosome activity in the fall. This may be associated with the organisms being better fed during this season and perhaps the storage of lipids by the pteropods for overwintering. It is known that other species of pteropods, including the closely related *Limacina helicina*, can store lipids as diacylglycerol ethers or triacylglycerols (Falk-Petersen et al., 2001; Gannefors et al., 2005). Odd-and long-chain fatty acid storage has also recently been demonstrated in *L. retroversa* (Boissonnot et al., 2019). This has been posited to be a tactic for coping with the strong seasonality in food supply that is prevalent in polar and boreal conditions. Due to the linkage between food availability and response to stress, explicit analysis of energetic stores and associated enzyme activity in the period leading up to the highest local CO_2_ conditions may provide important insights as we seek to understand the resilience of these populations to a changing GoM environment. As our in situ sampling for gene expression focused on the largest animals, it may also be important to look at the expression across size classes.

### 4.3 Biomineralization

Our shell condition and gene expression results support our hypothesis that annual changes in local carbonate chemistry influence the biomineralization of the local population of *L. retroversa*. The significantly lower shell condition that was observed in January occurred at an oversaturated state (Ω_Ar_ = 1.5), indicating that these organisms are influenced by saturation states above the dissolution threshold. These in situ observation of changes in shell condition have not been previously reported for field samples from the East Coast of North America, but are complementary to findings in Pacific upwelling and polar regions of the sensitivity of pteropod biomineralization to changing local environmental conditions (Bednarsek & Ohman, 2015; Bednaršek et al., 2012b) and recent observations near CO_2_ vent sites in the Mediterranean (Manno et al., 2019). This suggests that shell condition, assessed using a very simple stereomicroscope method, could be used as a good indicator of the biological consequences of changing saturation state, similar to the proposed use of pteropods for this purpose on the West Coast of the United States (Bednaršek et al., 2017; Weisberg et al., 2016). Thecosome pteropods are likely more sensitive to OA than many economically relevant species such as fish and crustaceans (Kroeker et al., 2013; Kroeker et al., 2010), and it appears that these changes in shell condition occur at saturation states that do not substantially influence the metabolism of the population, so they may serve as an early warning system for perturbations in the ecosystem, providing opportunities to take remedial action before tipping points are crossed for the broader community (Bednarsek et al., in review).

The observed patterns in shell condition are complemented by gene expression changes. Interestingly, patterns in biomineralization gene expression also appear to be influenced by life cycle, with transcripts associated with the extracellular matrix, collagen, and chitin synthesis and breakdown being upregulated in the fall during what appears to be a phase of growth. Collagen and chitin are thought to be important non-crystalline components of molluscan shell and have been shown to be associated with shell deposition (Zhang et al. 2012). They do, however, also have other biological roles, making definitive assessment of the physiological implications of our findings difficult. Analysis of GO terms indicated that gene expression in all other seasons was enriched with chitin binding and metabolic processes when compared with the fall. Biomineralization associated gene expression in January was, however, distinct from all other seasons. During this period of lower saturation state overall gene expression was similar to that of pteropods exposed in the laboratory to increased CO_2_ (Maas et al., 2018): during both January and the CO_2_ exposure experiments genes that are thought to be directly associated with calcification, including alkaline phosphatase and a carbonic anhydrase precursor, were upregulated. This suggests that there may be active calcification in the winter as the organisms work to repair environmentally damaged shells.

## 5. Conclusions

Our results show that seasonality in food availability, reproduction and carbonate chemistry interact are associated with variation in the population abundance, size distribution, physiology, and shell condition of pteropods in the Gulf of Maine. Populations show a consistent daytime vertical distribution in the upper ∼150 m with the largest portion of the community in the upper 50 m. Although patchy horizontal abundances make assessment of cohort development difficult, there appears to be a large reproductive event after the spring bloom followed by continued reproduction in the summer and fall. This supports the hypothesis that there are two major cohorts per year with different growth rates and spawning times. There is evidence supporting the idea that populations are supported by influx of individuals from offshore and are locally-reproducing. Whether individuals are adapted to the region’s seasonal carbonate chemistry patterns remains unclear, although there is evidence of changing physiology over the annual cycle that may be associated with seasonal acclimation to food and carbonate chemistry conditions.

Currently the periods of high food availability, pH, and saturation state are phenologically in sync with periods of reproduction, allowing the most sensitive early life stages to have apparent high success in the region. During the winter, when saturation states are lowest, there are gene expression and shell condition changes reminiscent of laboratory exposure to enhanced CO_2_, indicating that as anthropogenic CO_2_ continues to lower saturation states, pteropod populations may be locally impacted. We expect that the initial detrimental effects will be noticed during reproductive events in the fall. As it appears that there is active lipid storage and reproduction during this phase, any decrease in saturation state during this annual period may influence the success of overwintering pteropods in the region and serve as a population bottleneck. As pteropods are known to locally be prey for planktivorous fishes (Bigelow, 1924; Suca et al., 2018), and to play a role in modifying the alkalinity budget and carbonate chemistry in near bottom basins in the region (Wang et al., 2017b), assessing their abundance should remain a component of local monitoring, while additional shell condition assessment may be warranted.

## ACKNOWLEDGEMENTS

We would like to thank Captain K. Houtler and Mate I. Hanley for their support aboard the R/V *Tioga*. At sea sampling was supported by Phil Alatalo, Leocadio Blanco Bercial, Sophie Chu, Taylor Crawford, Maja Edenius, Katherine Hoering, Ian Jones, Robert Levine, Mike Lowe, Camille Pagniello, Lenna Quackenbush, Andrea Schlunk, Ali Thabet, and Peter Wiebe. We appreciate all the hard work and fantastic note taking of Nancy Copley and the analytical expertise of Katherine Hoering in the lab.

## DATA AVAILABILITY

Raw gene expression sequences are archived at NCBI (NCBI BioProject PRJNA260534).

---

Supplementary Data 1: Functional annotation (.xls)

Supplementary Data 2: Mapping success (.xls)

Supplementary Data 3: Differentially expressed genes, annotated (.xls)

Supplementary Data 4: GO Enrichment Analysis (.xls)

Supplementary Data 5: Dendograms and heatmaps of functionally important gene expression patterns (.pdf)

