## Supplemental Data 5 for "Seasonal variation in physiology and shell condition of the pteropod *Limacina retroversa* in the Gulf of Maine relative to life cycle and carbonate chemistry"

Dendrogram of TMM normalized FPKM values for genes from in situ samples that were identical to those that had been shown to be DE in laboratory $CO_2$ exposures (Maas et al. 2018). Each gene was normalized ((x-xmin)/(xmax-xmin)) and a dendrogram was created using the heatmap.plus function in R (calculated using euclidean distances and clustered by complete linkage; Blue = value of 0, Red = value of 1). Clusters were tested for statistical significance ($p < 0.05$, colored stars) using the Primer v7 statistical package using a Simprof test with 999 permutations.

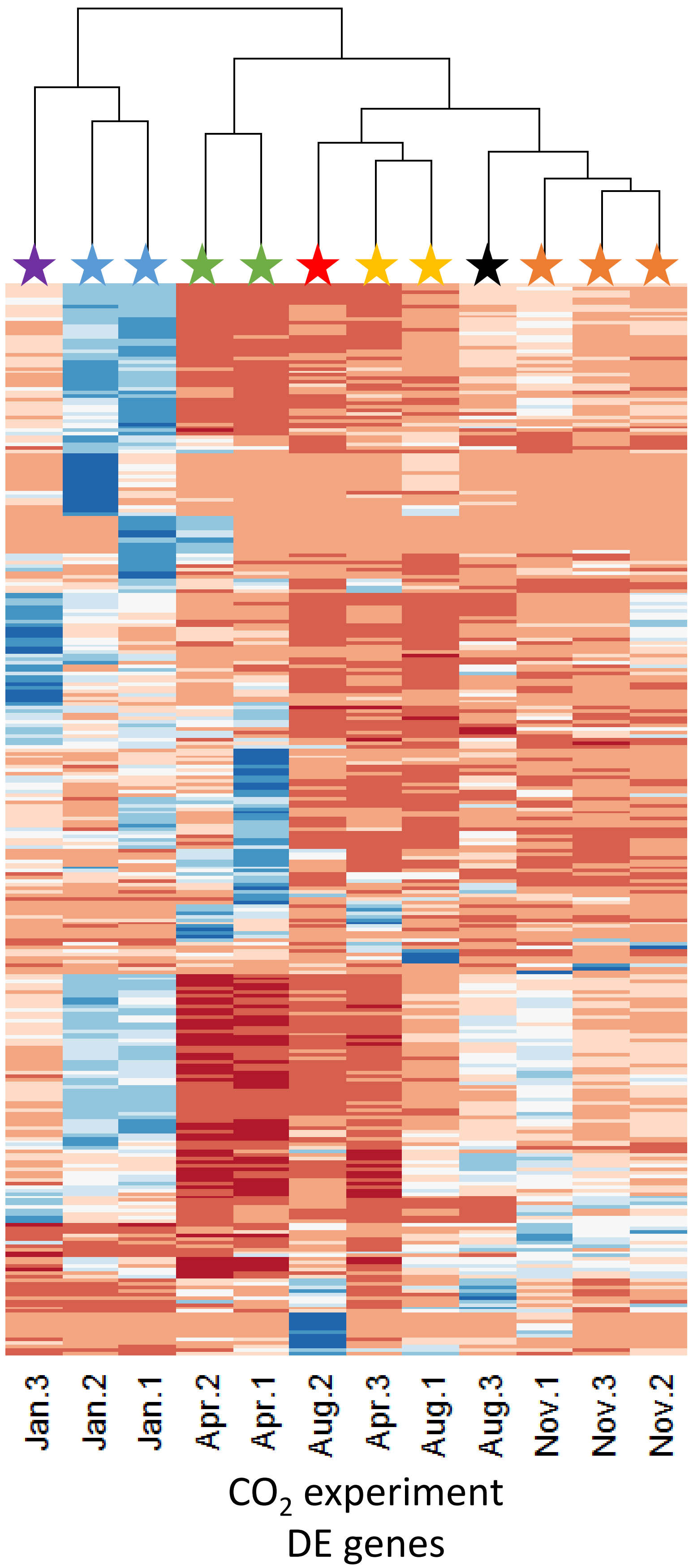

Dendrogram of TMM normalized FPKM values for genes from in situ samples that were identified as being associated with biomineralization (Zhang et al. 2012) or reproductive (Boutet et al., 2008; Ciocan et al., 2011; Tong et al., 2015) processes. Each gene was normalized ((x-xmin)/(xmax-xmin)) and a dendrogram was created using the heatmap.plus function in R (calculated using euclidean distances and clustered by complete linkage; Blue = value of 0, Red = value of 1). Clusters were tested for statistical significance (p <0.05, colored stars) using the Primer v7 statistical package using a Simprof test with 999 permutations.

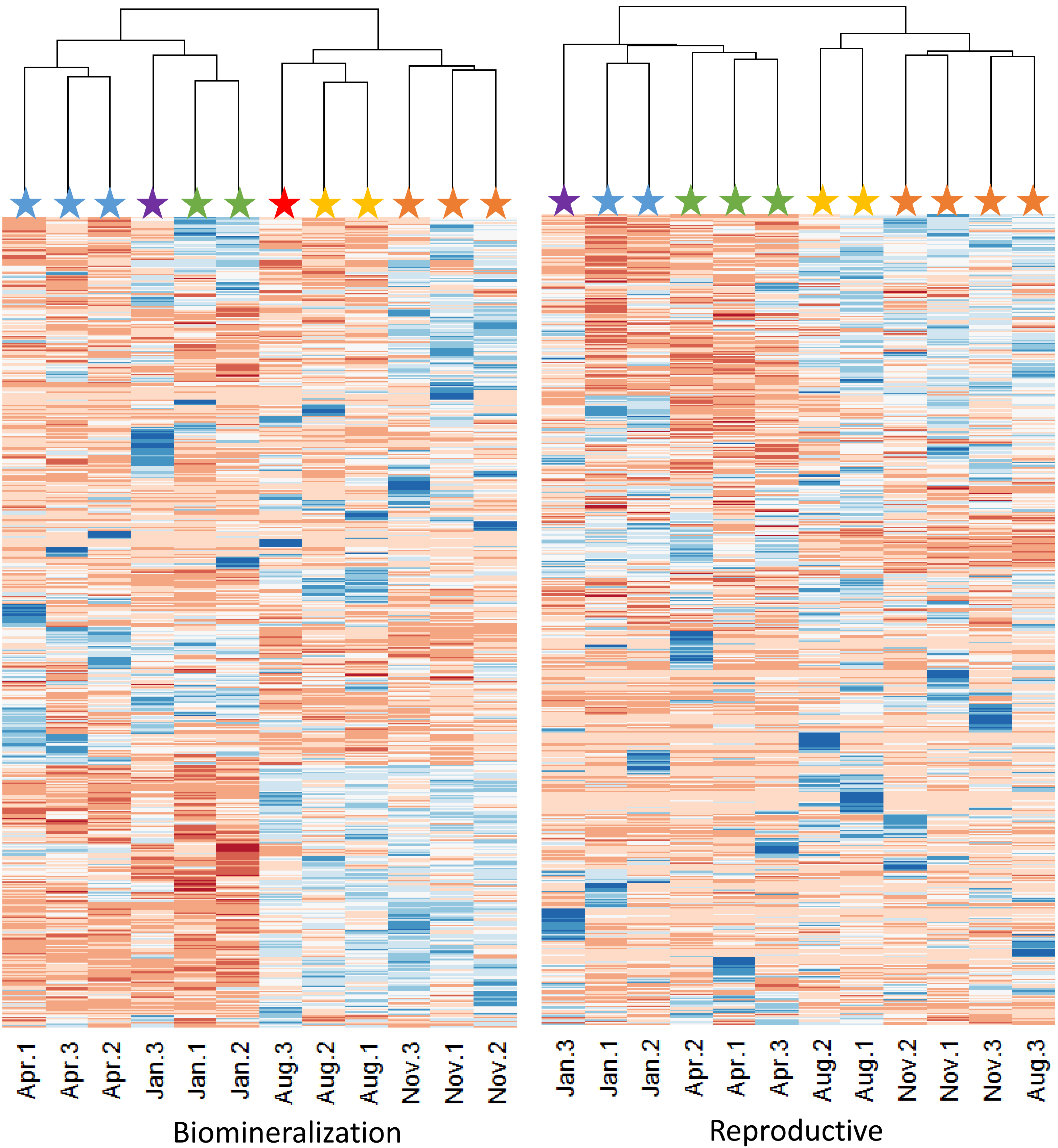

Dendrogram of TMM normalized FPKM values for genes from in situ samples that were GO annotated as having a metabolic or lipid function. Each gene was normalized ((x-xmin)/(xmax-xmin)) and a dendrogram was created using the heatmap.plus function in R (calculated using euclidean distances and clustered by complete linkage; Blue = value of 0, Red = value of 1). Clusters were tested for statistical significance (p <0.05, colored stars) using the Primer v7 statistical package using a Simprof test with 999 permutations.

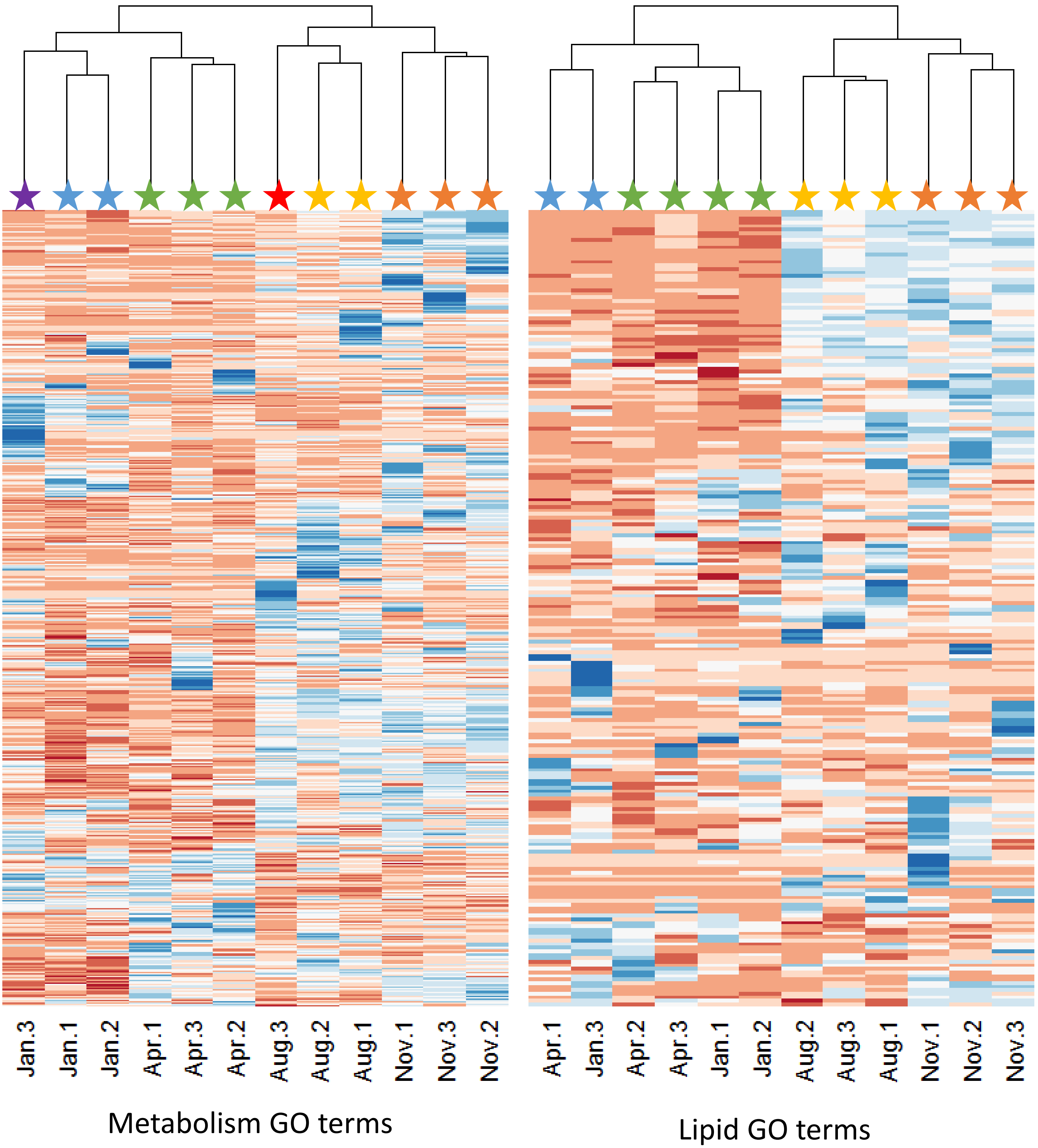
